# A scalable algorithm for clonal reconstruction from sparse time course genomic sequencing data

**DOI:** 10.1101/2021.08.19.457037

**Authors:** Wazim Mohammed Ismail, Haixu Tang

## Abstract

Long-term evolution experiments (LTEEs) reveal the dynamics of clonal compositions in an evolving bacterial population over time. Accurately inferring the haplotypes - the set of mutations that identify each clone, as well as the clonal frequencies and evolutionary history in a bacterial population is useful for the characterization of the evolutionary pressure on multiple correlated mutations instead of that on individual mutations. Here, we study the computational problem of reconstructing the haplotypes of bacterial clones from the *variant allele frequencies* (VAFs) observed during a time course in a LTEE. Previously, we formulated the problem using a maximum likelihood approach under the assumption that mutations occur spontaneously, and thus the likelihood of a mutation occurring in a specific clone is proportional to the frequency of the clone in the population when the mutation occurs. We also developed several heuristic greedy algorithms to solve the problem, which were shown to report accurate results of clonal reconstruction on simulated and real time course genomic sequencing data in LTEE. However, these algorithms are too slow to handle *sparse* time course data when the number of novel mutations occurring during the time course are much greater than the number of time points sampled. In this paper, we present a novel scalable algorithm for clonal reconstruction from sparse time course data. We employed a statistical method to estimate the sampling variance of VAFs derived from low coverage sequencing data and incorporated it into the maximum likelihood framework for clonal reconstruction on noisy sequencing data. We implemented the algorithm (named ClonalTREE2) and tested it using simulated and real sparse time course genomic sequencing data. The results showed that the algorithm was fast and achieved near-optimal accuracy under the maximum likelihood framework for the time course data involving hundreds of novel mutations at each time point. The source code of ClonalTREE2 is available at https://github.com/COL-IU/ClonalTREE2.

## 1 Introduction

Long-term evolution experiments (LTEE) are often used to study the mechanisms of adaptation and evolution [7] and for detecting genetic elements under selection [2] in clonal populations of unicellular microbes (e.g., *E. coli* [9, 23, 1, 19, 2]) as well as multi-cellular model animals (e.g., fruitfly [5]). The characterization of clones resulting from novel mutations in the population and their ancestral relationship is a critical first step for subsequent evolutionary and functional analyses. A straightforward approach to achieving this goal is to random sample and sequence a large number of individual clones from the population. In practice, however, this approach is quite expensive even with the reduced cost of DNA sequencing. As a result, *bulk sequencing* is often adopted, in which the variations and the *variant allele frequencies* are derived from the whole genome sequencing (WGS) of mixture of clones randomly sampled at different time points during the evolution process.

It is non-trivial to reconstruct the clonal structure from the variant allele frequencies (VAF) [14]. Previously, we formulated the *time course clonal reconstruction problem* using a maximum likelihood framework [15], and proposed several heuristic greedy algorithms to solve the problem. In addition, we extended the algorithms to addressing the *sparse* time course data, which were acquired from the number of time points fewer than the number of novel mutations. We evaluated these algorithms using simulated experiments and observed that the greedy algorithms could accurately reconstruct as many clones as those reconstructed by a brute-force algorithm or even better on average, while improving significantly on speed. In particular, we devised the GP-GT algorithm that quickly and accurately reconstructed clones for clonal populations containing 30 or fewer novel mutations. Therefore, it achieved satisfactory results on the data from the LTEE of wild-type *E. coli* [2].

Despite the success, the greedy approach has some limitations. First, it assumes the input VAFs are the true allele frequencies in the population, while in real cases, the VAFs are approximate statistical estimates of the true allele frequencies with a positive sampling variance depending on the number of individuals sampled from the population as well as the read coverage of the sequencing data [12]. This noise in allele frequency estimation can drastically reduce the solution space of clonal trees searched by our algorithms, and in turn reduce the chances of finding the true solution.

More importantly, the algorithm is sufficiently fast only when the number of novel mutations in the population during the period of observation is relatively small (below 30). The strains studied in the LTEE of *E. coli* [2] include wild-type and mismatch repair (MMR) deficient strains. The number of mutations expected to be observed in three years of evolution (≈ 2500 generations at 1 ml transfer density) is approximately 2.5 with a wild-type mutation rate of 1 × 10^−3^ [8]. For this case, the running time of the *GP-GT* algorithm is quite reasonable. However, for the MMR deficient strain of *E. coli*, the mutation rate is 150 times higher than that of the wild-type, and hence, we expect several hundred novel clones occur during the period of observation, which makes the time course data very sparse (the number of time points is only six). As a result, the greedy algorithms developed previously [15] are too slow to handle these sparse time course data.

In this paper, we present a novel scalable algorithm for clonal reconstruction from sparse time course data containing hundreds of novel mutations occurring at each sampled time point. We employed a statistical method to estimate the sampling variance of VAFs derived from low coverage sequencing data and incorporated it into the maximum likelihood framework for clonal reconstruction. We implemented the algorithm and tested it using both simulated and real sparse time-course genomic sequencing data. The results showed that the algorithm could finish within reasonable time while achieving satisfactory accuracy on sparse and noisy data.

## 2 Methods

### 2.1 Time Course Clonal Reconstruction Problem

As previously described [15] we model the problem using the clonal theory of evolution [24, 22] which assumes that all bacterial cells in an evolving population are descendants of a single *founding clone* forming new clones by accumulating mutations following the *infinite sites assumption*, i.e., a mutation occurs at a single locus at most once during the period of evolution experiment. The ancestral relationships between clones in such a model can be represented by a rooted, directed tree *T* known as the *clonal tree* where nodes represent clones and edges represent direct ancestral relationships. We define the *time course clonal reconstruction problem* (TCCRP) as the problem of inferring the unknown clonal tree T from a matrix of *variant allele frequencies* (VAFs) across all variational sites in a bacterial genome at a series of *n* time points [6], i.e., *F* = [*f_i,j_*], where *f_i,j_* indicates the allele frequency of the variant *j* at the time point *i*. Notably, each variant is first introduced by a mutation (or a set of mutations) *j* at the time point *t_j_*, generating a novel clone (denoted by the specific mutation *j*) from its parent. Note that, here, *t_j_* is the earliest time point *t*, such that *f_t,j_* > 0, and for ∀*t* < *t_j_, f_t,j_* = 0. Here we consider only single nucleotide variations but the same approaches can be easily extended to other kinds of genomic variations.

We formulated the TCCRP using a maximum likelihood framework [15]. Given the input matrix *F* = [*f_i,j_*], where 1 ≤ *i,j* ≤ *n* over *n* novel mutations (or novel clones) *sorted* over *n* time points (i.e., each mutation introduced at a known distinct time point), we want to find a directed tree *T*\* = {(pr(*i*), *i*)| *i* = 1, 2,…, *n*} on *n* nodes (where pr(*i*) is the *unique* parent node of node *i*), that maximizes the following likelihood function,

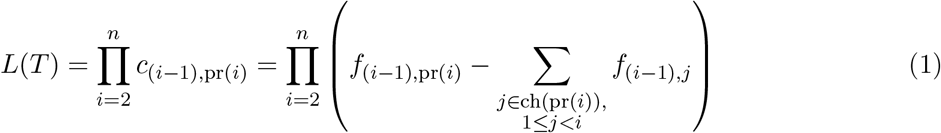

where *c_i,j_* represents the (unknown) frequency of the clone *j* at the time point *i*, and ch(*i*) represents the set of all children of the node *i*. The probability of introducing the mutation *i* in the clone pr(*i*) between the time points *i* — 1 and *i* is approximated by the frequency of the clone pr(*i*) at the time point *i* – 1, which can be computed using the relationship between the *clonal frequency matrix, C* = [*c_i,j_*], the *allele frequency matrix, F* and the *clonal tree, T* given by 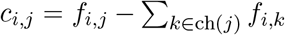 [6]. Note that a clonal tree becomes invalid if

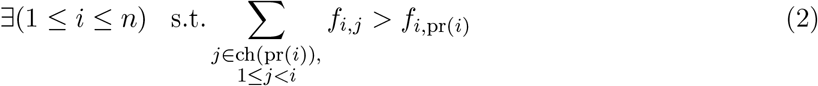

All elements of *C* must be non-negative and the rows of *C* must sum to one. We proposed a heuristic *greedy tree (GT)* search algorithm [15] to solve the TCCRP. The algorithm grows the directed tree from the root node (founder) such that at each iteration *i* > 1,

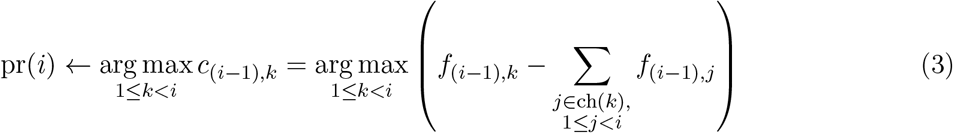

provided pr(*i*) does not lead to an invalid solution (equation 2). At any given iteration, if the optimal choice led to an invalid solution, a backtracking approach is used to pick the next optimal greedy choice until all valid solutions are explored. The worst case running time of this algorithm is *O*(*n*!), although in the best case it runs in *O*(*n*^2^) time.

In practice, because of the often scattered genomic sequencing conducted in a time course, we may observe many mutations occurring at the same time point *i*, i.e. F may not be a square matrix. In this case, if the occurrence order of these mutations is determined, we assume that the VAFs of all variants remain approximately constant between the time points *i* – 1 and *i*, and thus we can apply the GT algorithm to identify the maximum likelihood clonal tree. In order to search for the optimal order of *n_i_* mutations at a specific time point *i*, we employed a heuristic *greedy permutation (GP)* approach: all permutations of the *n_i_* mutations at time point *i* are examined at the iteration by the GT algorithm and the permutation producing the maximum likelihood clonal tree is chosen and fixed for the subsequent iterations. Therefore, the resulting GP-GT algorithm searches for the clonal tree for each of *n_i_*! permutations at the time point *i*. When *n_i_* is small (e.g, *n_i_* < 5), the GP-GT algorithm can achieve satisfactory results within reasonable time [15]. We showed that the temporal order of mutations that can be inferred from the time series allele frequency data reduces the solution space of all possible clonal trees. Note that our problem formulation differs from that of another related work - Calder [16], in terms of the maximum likelihood framework which provides a scoring scheme for ranking multiple valid solutions and thus formulates it as an optimization problem.

Unfortunately, the GP-GT algorithm encounters two challenges when applied to real-world time course genomic sequencing data from bacterial populations. First, the VAF estimates from sequencing data are not accurate; as a result, many clonal trees (including the true solution) in the solution space may become invalid according to equation 2. Second, when the number of mutations occurring during the LTEE is much greater than the time points from which the pooled sequencing data are acquired, the number of new mutations per time point (i.e., *n_i_*) is large (e.g., in the data from mismatch repair deficient *E. coli* populations [2], more than 50 mutations were observed at a single time point). In these cases, the GP algorithm needs to explore a huge space (*n_i_*!) of mutation permutations at each time point, and becomes too slow. Here, we present a scalable algorithm for clonal reconstruction from time course genomic sequencing data that addresses these two challenges.

### 2.2 Penalizing the likelihood score for invalid solutions caused by noise

The strict definition of validity given by equation 2 may classify many clonal trees, including the true solution to be invalid if the input VAF estimates are noisy. We define the problem of finding the optimal clonal tree from noisy VAF input as the noisy version of the TCCRP. To address this problem, we aim to remove the condition defined in equation 2 and replace it with a penalty score in the likelihood function (equation 1) for each solution producing negative elements in matrix *C*. This penalty should be proportional to the sum of all negative clonal frequencies resulting from that solution. We define the penalty function as described below. Firstly, we note that the minimum achievable standard deviation *σ_f_i,j__* of VAF estimate *f_i,j_* is given by [12],

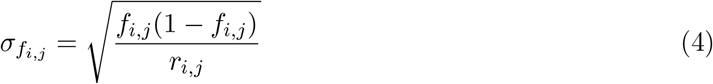

where *r_i,j_* is the depth of coverage at the locus corresponding to the variant *j* from the sample at time point *i*. We assume that the number of sampled individuals is high enough that the sampling variance depends only on the number of reads sampled from the pooled DNA and not on the number of sampled individuals. This is a reasonable assumption for microbial population sequencing where the number of individuals collected for bulk sequencing is usually large. For any given solution, for each *i, j* such that

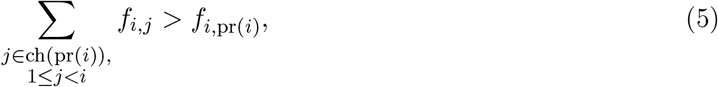

the penalty *p_i,j_* is defined as

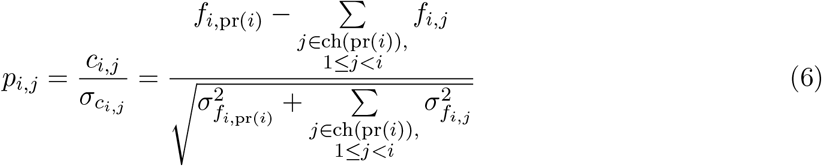

This penalty is then added to the likelihood score for that solution. The rationale for dividing the negative clonal frequencies by their corresponding standard deviations is that the estimated standard deviation gives us the variability of the clonal frequency due to noise and thus becomes an approximation of how likely it is for the clonal tree to produce a negative clonal frequency. So when the standard deviation is small but the magnitude of the negative clonal frequency is large, a higher penalty must be assigned to the solution compared to the solution in which the standard deviation is also large.

Note that this approach removes the need for backtracking steps in the greedy algorithms because no solution is considered invalid and the solution at any given iteration is chosen in a greedy manner based on the best likelihood score including the penalties for negative clonal frequencies during that iteration. This improves the speed of the algorithm compared to the backtracking approach as it converges to the optimal solution much faster while exploring a smaller number of solutions.

### 2.3 Reducing the permutation search space

The second challenge we attempt to address is the potentially large search space of mutation permutations (by the GP algorithm) due to the number of novel mutations (clones) being much larger than the number of time points. We first investigate the ancestry graph *G* [6] defined as the directed acyclic graph (DAG) constructed by comparing all pairs of columns of *F* and adding edges only if the ancestry condition is met. According to the *ancestry condition*, variation *v_s_* can be the ancestor of variation *v_t_* only when the VAF *f_i,s_* is greater than *f_i,t_* across all samples *i*. The ancestry condition is a necessary (but not sufficient) condition for a valid solution and the clonal tree is always a spanning arborescence of the ancestry graph.

As illustrated in Figure 1, we adopt the ancestry graph approach to characterizing the optimal permutation of a group of mutations occurring at the same time point (e.g., the red, green and yellow mutations in Figure 1A). We first construct an ancestry graph on the subset of mutations under consideration at a specific time point, which is an induced subgraph (referred to as the ancestry subgraph; Figure 1B) of the ancestry graph on all mutations (clones). The ancestry subgraph is also a directed acyclic graph that induces a partial order of the mutations. Any order that violates this partial order violates the ancestry condition and hence is invalid. As a result, significantly fewer number of permutations need to be explored.

**Figure 1:**
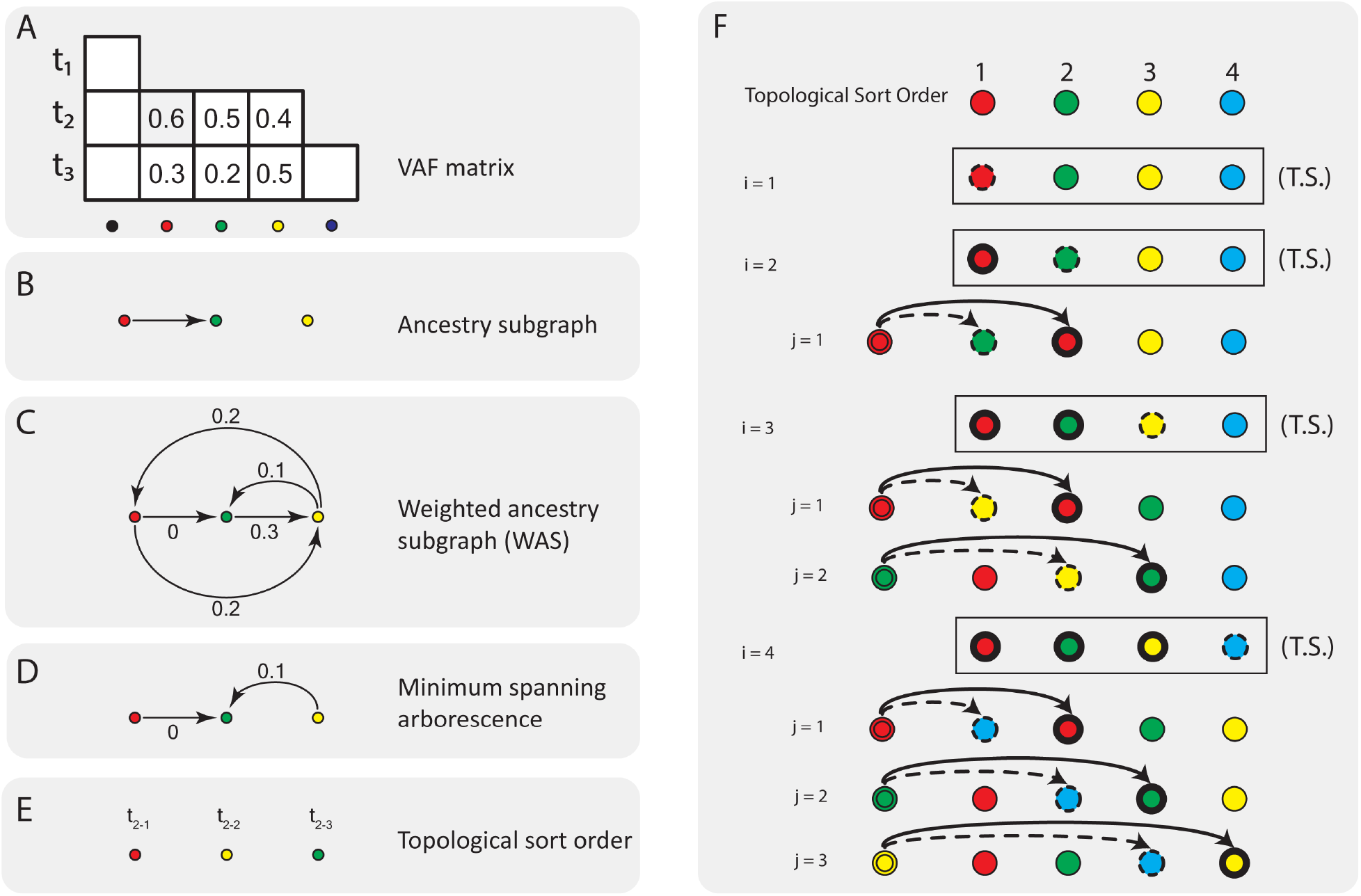
Illustration of reducing the combinatorial search space of permutations of unordered mutations. (a) A toy VAF matrix on three mutations, represented in red, greed and yellow, respectively, that appear to spawn at the same time point (t2). To obtain a partial order for these mutations, we first construct (b) the ancestry subgraph that shows all the putative ancestral relationships among these mutations that satisfy the ancestry condition. From that we construct (c) a *weighted ancestry subgraph (WAS)*, which, in addition to all edges in the ancestry subgraph, contains two edges between every pair of vertices not connected in the ancestry subgraph. The weights assigned to the edges represent the “errors due to noise” parameters that show the deviations from the ancestry condition if the respective edges are considered legitimate ancestral relationships. Minimizing the error due to noise can be achieved by finding (d) the minimum spanning arborescence of this graph, which provides a partial order given by (e) its topological sort order. (f) Given the topological sort of the mutations (top) of unordered mutations, we can enumerate their potential permutations through two nested loops: the outer loop (i) iterates over the mutations shown as dotted circles, and the inner loop (*j*) iterates over mutations shown as bold circles. For each inner loop *j*, the mutation *v_i_* is placed in front of *v_j_*, assigning *v_j_* the same ancestor as that of Vi shown here as a double circle that matches the same color as *v_i_*.

Note that when we construct the ancestry subgraph from a noisy VAF matrix *F*, the frequencies of potential ancestors may be lower than the total frequencies of their children in some samples due to noise. So this graph will result in some missing edges resulting in more number of partial orders to be explored than necessary. To circumvent this problem, we construct a weighted ancestry subgraph (WAS) that models the errors due to noise. First, all the edges in the ancestry subgraph are assigned to zero weights. In addition to these edges, we add two edges one in either direction between every pair of mutations, *v_s_* and *v_t_* where not all samples *i* have frequencies *F_i,s_* > *F_i,t_* or *F_i,t_* > *F_i,s_* (Figure 1C). The edge weights are assigned as,

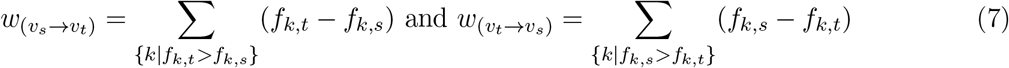

Figure 1C shows the WAS constructed from the partial allele frequency matrix *F* shown in figure 1A. We assume that the edge weights represent the likely error due to noise, i.e., the deviations from the ancestry condition if the respective edges are considered as legitimate ancestral relationships. We aim to minimize the total error due to noise by finding a minimum spanning arborescence of the WAS. We note that the WAS is not a DAG, and thus it does not induce a partial order of mutations. But the minimum spanning arborescence of the WAS is a DAG, and induces a partial order. We will exploit this order as our guide to reduce the search space of permutations.

Note that even when we obtain a partial order of mutations, there might exists too many valid permutations over hundreds of mutations occurring at the same time point. A critical observation is that the likelihood score of a clonal tree is dependent on the order of two mutations only if these two mutations share a common ancestor. Following this observation, the clonal tree reconstruction algorithm needs to search for the optimal permutation of unordered mutations only in a particular subset of permutations depending on the previously determined ancestors. Specifically, for each set of unordered mutations, we start with a topological sort order following the minimum spanning arborescence of the WAS constructed from the allele frequency matrix *F* (Figure 1F). For each mutation *v_i_* (dotted circles in figure 1F) in this set, we switch its position to the left of each mutation *v_j_* (bold circles) before *v_i_* in the topological sort order, and assign the same ancestor of *v_j_* (double circles) to *v_i_* while keeping the rest of the ancestral relationships the same as the optimal clonal tree. We then re-compute the likelihood of the resulting clonal tree. If the likelihood of the new clonal tree is higher than that of the optimal clonal tree, we replace it with the new tree. Finally, the clonal tree with the best likelihood score is output. This algorithm (ClonalTREE2: Algorithm 1) runs in *O*(*n*^2^) time on n unordered mutations occurring at the same time point, which is significantly faster than the GP algorithm that runs in *O*(*n*!) time.

## 3 Results

### 3.1 Simulation Experiments

We followed the previously published simulation procedure [15] with a few changes as follows. In order to truly mimic experimental LTEE data, we applied the same quality control filters to the simulated data - 1) all allele frequencies *f_i,j_* < 0.05 were set to 0; 2) all variants (columns in *F*) without non-zero frequencies in at least two time points are removed, and we assigned the clonal frequencies at each time point by sampling a Dirichlet distribution with parameters given by the clonal frequencies at the previous time point instead of a uniform random distribution as used previously. In addition to the simulation of the accurate allele frequency data, we simulated noisy allele frequency data as follows. For each simulated allele frequency *F_i,j_*, we simulated the depth of read coverage *r_i,j_* sampled from a normal distribution with the mean of 170 and the standard deviation of 40. These parameters were chosen to be close to those of the LTEE experiment [2]. Next, using this depth of coverage, we computed the standard deviation *σf_i,j_* of the allele frequency *f_i,j_* using equation 4. Finally, we generated a corresponding noisy allele frequency *f_j_* by sampling from a normal distribution with mean *f_i,j_* and standard deviation *σ_f_i,j__*. We simulated 100 datasets with different number of clones - 20, 30, 40, 50, 100, 200, 400, 700 and 1000, respectiely. We generated both the accurate and noisy allele frequencies data in each simulation experiment. The number of time points in these datasets is sampled from a binomial distribution *B*(*n*, 0.6), where n is the number of clones in the respective simulation experiment.

#### Algorithm 1

ClonalTREE2 algorithm

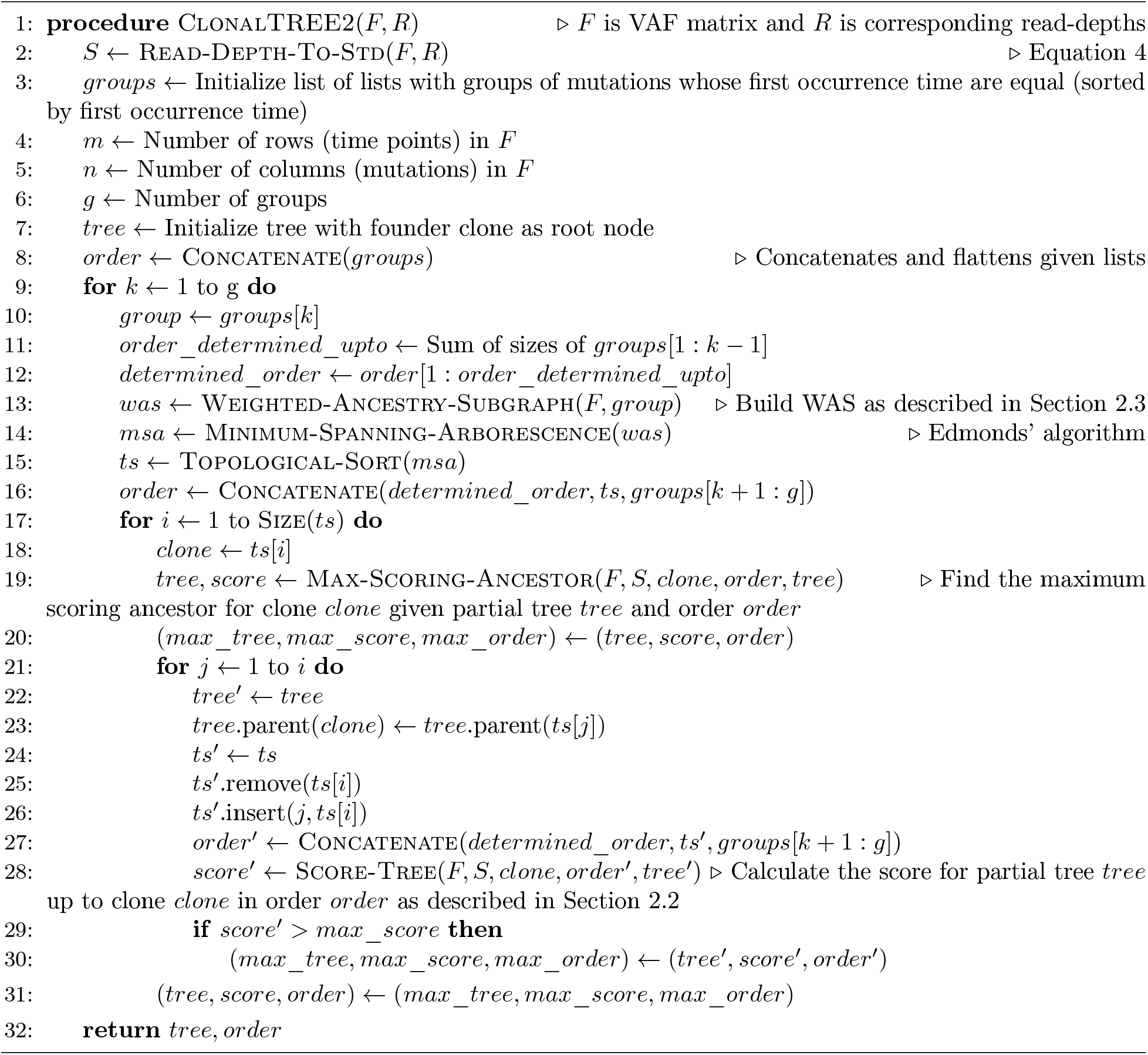

We compared the performance of Calder [16], ClonalTREE (i.e., the GP-GT algorithm published previously [15]), ClonalTREE_P (the revised GP-GT algorithm that incorporates the penalty function in the likelihood optimization) and ClonalTREE2 on both the accurate and noisy allele frequencies data in terms of their *recall* (defined as the proportion of clones completely and correctly reconstructed by the algorithm), *running time* and *clonal accuracy* (defined as the proportion of variants that are correctly assigned to all clones) (Figure 2). We used the default parameters and the Gurobi ILP solver when running Calder. Because Calder was designed for diploid genomes, we multiplied allele frequencies from our simulations by 0.5 before input to Calder. We also applied the same set of quality control filters before running all of the algorithms so that the number of variations input to each algorithm is consistent.

**Figure 2:**
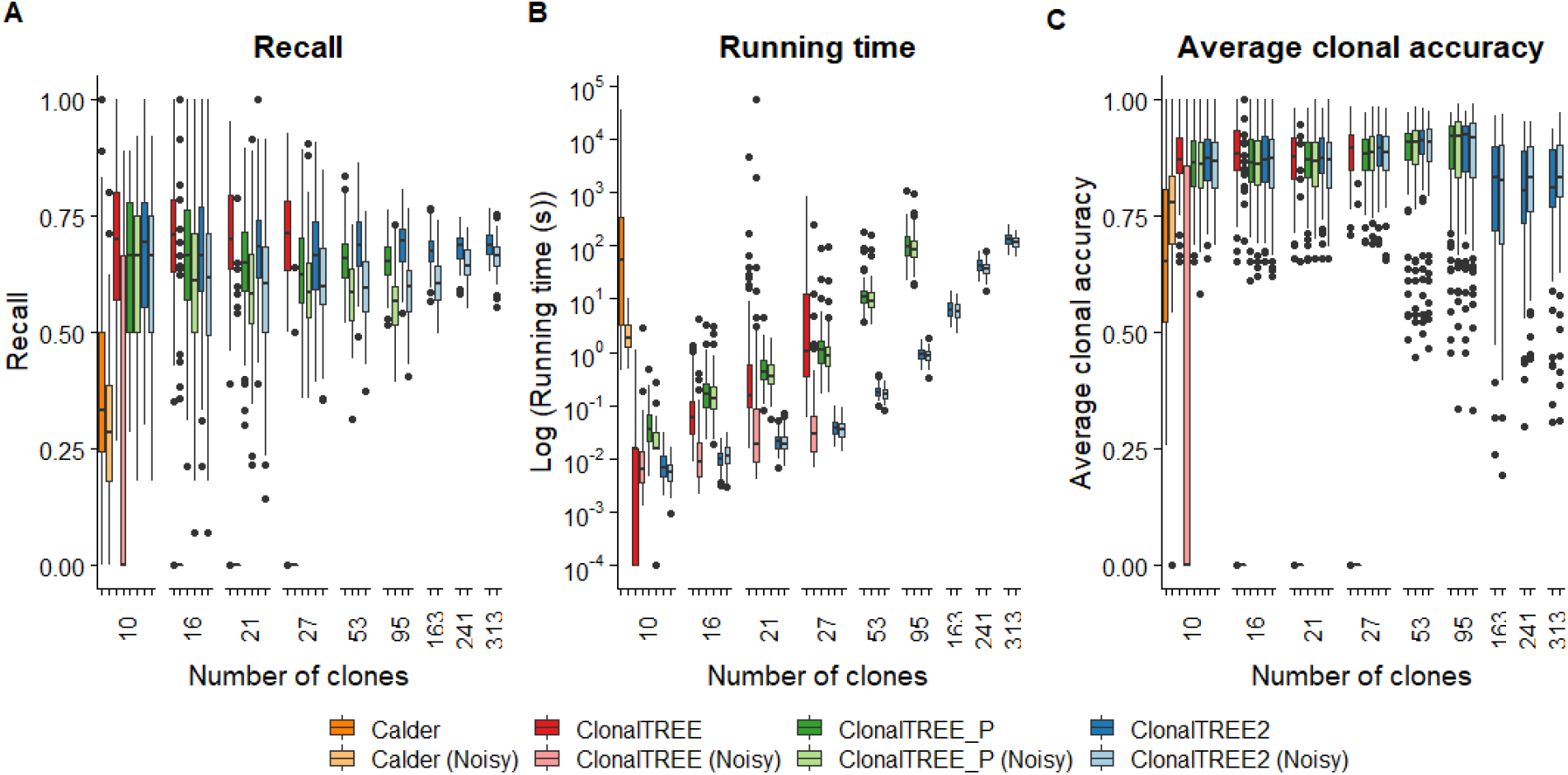
Comparison of the recall, running times and average clonal accuracy of Calder, ClonalTREE, ClonalTREE_P and ClonalTREE2 on the accurate and noisy allele frequencies for different number of clones in the simulation experiments. The results are computed over 100 simulations with different number of clones - 20, 30, 40, 50, 100, 200, 400, 700 and 1000 clones, respectively. The x-axis shows the average numbers of remaining clones across 100 simulations after applying the filters.

The results of the simulation experiments showed that while the performance of ClonalTREE on the accurate data achieved about an average of 70% recall, and 85% average clonal accuracy in most accurate datasets, it does not perform well on noisy data. This is because the number of invalid solutions are high when the data is noisy and in many cases the algorithm fails to find any solution. The incorporation of the penalty score significantly improves the performance of ClonalTREE_P on noisy data because in this algorithm every solution is taken into consideration while the likelihood score with the penalty score provides a gradient among all solutions. The downside of ClonalTREE_P is that it is very slow on the experiments involving more than 30 clones. The running time of the algorithm increases from a few minutes to several hours depending on the number of unordered mutations at each time point (*O*(*n*!)). Hence, It is not practical for the LTEE data containing more than 30 mutations (e.g., for the MMR-deficient E. coli strains). ClonalTREE2 runs much faster than ClonalTREE and ClonalTREE_P, while its recall is almost as good as or better than the other algorithms on average: the average recall is close to 70% and the average clonal accuracy approaches 80% in all datasets, and it only spends less then five minutes to compute the clonal tree involving more than 300 novel mutations occurring at the same time point.

We observe that Calder runs much slower than ClonalTREE, with the ILP step specifically contributing to the majority of running time. We ran Calder only on the first dataset because the program took longer than 10 hours to complete for each simulation with number of clones more than 15. Similarly, we ran ClonalTREE only on the datasets with less than 30 clones, and ClonalTREE_P only on the datasets with less than 100 clones because their running times were too long for the datasets with more clones. Calder produces an average recall of about 35%, which is much lower than ClonalTREE2 on these simulations. It should be noted that ClonalTREE2 is designed to find the maximum likelihood solution provided the likelihood assumption is true in the LTEE. Since the simulation algorithm generates data that is consistent with the likelihood assumption, ClonalTREE2 is bound to find better solutions compared to Calder, as Calder was not designed to find the maximum likelihood solution. In cases where the likelihood assumption does not hold, the solutions provided by Calder might be closer to the true clonal tree.

## 4 Clonal reconstruction on the *E. coli* populations

We applied ClonalTREE2 to the time course genomic sequencing data from the *E. coli* LTEE [2], in which 50 lines of wild-type and 50 lines of mismatch repair deficient (MMR-) *E. coli* populations were maintained and observed for over 3 years. Of these 100 populations, ten populations (population ids: 113, 125, 126, 129, 210, 221, 233, 326, 410 and 417) were sequenced for every six months, resulting in six time points of pooled Illumina sequencing reads. At the end of three years, eight clones from each population were isolated and sequenced. Three of the ten populations are wild-type (populations 125, 129 and 221) and seven are MMR-strains (populations 113, 126, 210, 233, 326, 410 and 417). We used ClonalTREE2 to reconstruct the clonal evolutionary history for each of these 10 populations.

We detected the mutations by mapping the reads to the reference genome *E. coli* K-12 MG1655 [3], and estimated the allele frequencies of the variations in the *E. coli* population sampled at the six time points, using the bioinformatics pipeline described previously [15]. Briefly, we first removed adapters and low quality bases using Trimmomatic version 0.33 [4], then mapped the reads to the reference genome using bwa-mem version 0.7.12 [11]. We applied the following quality control filters to remove unreliable base calls. We first removed reads supporting bases with forward/reverse read balance less than 0.25. Then we called variants only when all of the following conditions are satisfied: 1) the VAF, estimated by the ratio of the number of reads supporting the variant allele to the total number of reads covering the site (supporting both the variant and the reference allele) is above 0.05, 2) the total number of reads at the site is above 10; and 3) the number of reads supporting the variant allele is above 6. We removed the variants without non-zero frequencies at two or more consecutive time-points, and the variants whose estimated allele frequencies were greater than 0.95 in four or more time points, or greater than 0.9 in all six time points. These variants are likely to be present in the founder clone instead of novel mutations forming novel clones. We considered only single nucleotide variations (SNV) in this analysis. We identified 14, 18 and 10 mutations from the wild-type populations and 420, 10, 314, 72, 555, 115 and 899 mutations from the MMR-deficient strains respectively. As expected, we observed that the number of variations in the wild-type populations are much smaller compared to those in the MMR-deficient strains (with the exception of the population 126 which is low due to a low read coverage in sequencing data).

To evaluate the accuracy of the reconstructed clones by ClonalTREE2, we compared the reconstructed clones with the eight clones that were isolated and sequenced at the end of three years of LTEE. The clonal compositions (haplotypes) of the sequenced clones were taken from Behringer et al. [2]. We compared only those variants in each sequenced clone that were also detected in the time series sequencing data and thus were used in clonal reconstruction. This set of variants is considered as the *known clonal composition (KC)* for each sequenced clone, against which we compared the *predicted composition (PC)* of each reconstructed clone. For each sequenced clone, we calculated the number of variants that are common in the known composition and the predicted composition (Figure 3). Note that the number of sequenced clones shown here is less than eight in some populations because we removed duplicated clones that share identical compositions of variants. Population 125 shows satisfactory clonal accuracy (85-100%). In the populations 129 and 221, the number of variants shared by the sequenced clones and those observed in time course genomic sequencing are small, and thus were not useful to evaluate the prediction performance by ClonalTREE2. In the populations 113 and 210, the accuracy of the reconstructed clones is satisfactory (59-97%), while in the population 326, the clonal accuracy is relatively low (33-47%), since the number of variants is very large compared to the number of time points, which significantly decreases the chance that the true clonal tree reflecting the mutations in the population is the one with the maximum likelihood that the ClonalTREE2 algorithm seeks. On the other hand, in the populations 233, 410 and 417, the number of variants shared by the sequenced clones and those observed in time course genomic sequencing are too small. Overall, in a majority of populations, ClonalTREE2 reconstructed the clones with satisfactory accuracy, comparable with the accuracy in the simulation experiments. These results indicate ClonalTREE2 can be used for clonal reconstruction from real time course genomic sequencing data. The predicted clonal tree can be used to compute clonal frequencies over time, which provides useful insights about selections on the novel clones.

**Figure 3:**
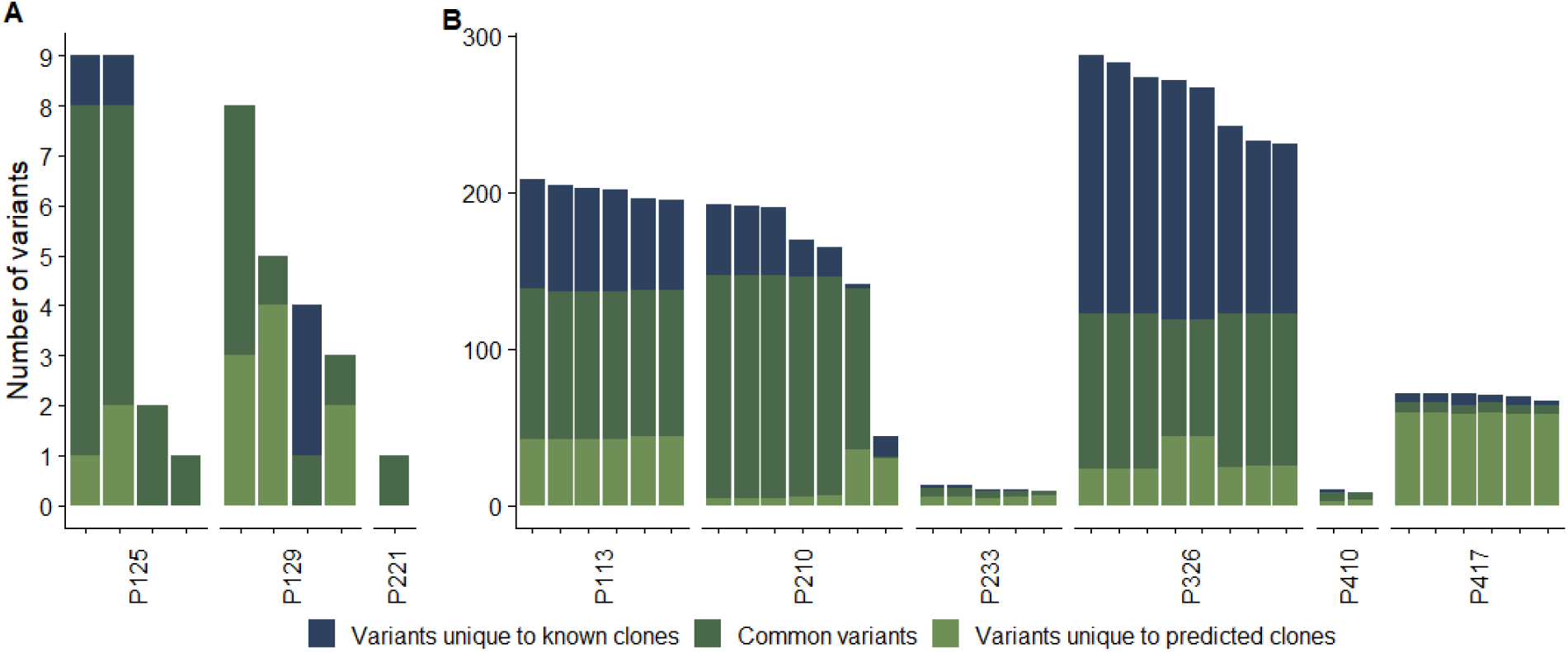
Comparison of variants inferred from the clonal sequencing and reconstructed clones using ClonalTREE2 in wild-type (left) and MMR^−^ (right) *E. coli* populations. Each bar represents a single pair of known/predicted clones.

## 5 Discussion

Algorithmic approaches to characterizing the clonal evolutionary history of an evolving cell population is critical not only for understanding the mechanisms of evolution in microbes and microbial communities ([2, 9, 10, 17, 20, 25]), but also for addressing important problems in cancer genomics, e.g., for reconstructing tumor genomes and predicting putative driver mutations ([18, 21]). The algorithmic problem of clonal construction demonstrates several combinatorial and probabilistic nature of the problem [22, 13, 6, 14], indicating some useful avenues to tackle the computational challenges presented. In this paper, we present ClonalTREE2, a scalable algorithm for clonal reconstruction from time course genomic sequencing data under a maximum likelihood framework. The algorithm addresses two specific challenges - 1) sampling noise due to shallow sequencing depth, and 2) data sparsity due to a large number of mutations occurring in the population. Through simulation experiments that resemble the LTEE, we showed that ClonalTREE2 overcomes these challenges without compromising the prediction accuracy.

## Acknowledgements

This work is partially supported by a Multidisciplinary University Research Initiative Award W911NF-09-1-0444 from the US Army Research Office, the National Institute of Health grant 1R01AI143254), and the Indiana University (IU) Precision Health Initiative (PHI).

